# Virtual-cell verification enables self-auditing AI discovery for immune rejuvenation

**DOI:** 10.64898/2026.08.04.742916

**Authors:** Yue You, Xueying Fan, Guangpeng Li, Wenchan Deng, Yunlin Fu, Huimin Hu, Wenle Ren, Shihong Lu, Guobin Han, Jian Shao, Shuangjia Zheng, Kaixin Zhou, Jiaming Kong, Jizheng Chen, Xiaodong Liu, Luyi Tian

## Abstract

Artificial-intelligence agents propose drug-discovery hypotheses faster than experiments can test them, yet their conclusions are rarely verified—against the underlying biology, the predicted perturbation, or the agent’s own scoring logic. We close this verification gap with an agentic framework built on three verifiers. First, PACE, a phenotype verifier, resolves immune aging into ten directionally scored, cell-type-resolved gene-set modules, selected for cross-cohort stability across four PBMC cohorts, and outperforms five established aging clocks in an independent in-house aging cohort of 434 elderly donors. Second, CellQ, a virtual-cell verifier built with multi-modal LLM, compresses each single-cell transcriptome into eight discrete tokens aligned to a language model’s vocabulary through residual vector quantization; it attains state-of-the-art perturbation prediction and uniquely resolves the weak, module-level shifts that differential-expression recovery misses. Third, an Analyzer–Planner–Auditor agent verifies its own scoring logic: screening 110 compounds in primary human PBMCs, it found aged-down modules more reversible than aged-up modules and revised its objective from an equal-weight mean to a balance-constrained minimum—a self-correction that generalized to an independent 13-compound T-cell assay. By verifying its predictions and its own objective against experiment, the framework points beyond hypothesis-generating AI toward self-correcting AI scientists whose objectives could continuously evolve.

## Introduction

Transcriptome-guided drug discovery prioritizes compounds by the cellular state they induce [1, 2], and has produced increasingly capable platforms for both repurposing and *de novo* design—structure-to-transcriptome virtual screening, exemplified by GPS [3], and signature-guided prioritization with lab-in-the-loop refinement, exemplified by DrugReflector [4]; in aging specifically, transcriptomic signatures of youthful cell states have been used to nominate candidate geroprotectors [5]. These approaches share a premise: that the therapeutic goal can be written as a single, coherent target signature known in advance. Many of the complex phenotypes and diseases that matter most violate this premise. Therapeutic benefit frequently arises from distributed, multi-target action rather than the modulation of a single axis—diseases are typically polygenic, cellular networks are buffered by redundancy, and most approved drugs were discovered through their net phenotypic effect rather than against a predefined target [4]. Immune aging is a paradigm case [6]: large single-cell atlases, building on earlier bulk-blood transcriptomic profiling [7], show that age-associated change is dispersed across cell types and partially coordinated gene programmes rather than concentrated in one canonical signature [8, 9], and existing single-cell immune-aging clocks (sc-ImmuAging [10], hUSI [11], SenCID [12])—which extend the foundational DNA-methylation clock tradition [13–16] to transcriptomics—compress this structure into useful scalar estimates that cannot resolve which programme moves, in what direction, or in which cell type under perturbation. For such phenotypes the central difficulty is not reconstructing a known signature but verification: determining whether a candidate actually shifts a set of weak, directional, cell-type-resolved phenotype axes, and whether the rule used to aggregate those axes into a single score remains valid once experimental data arrive.

Computational prediction of perturbation responses is the natural instrument for such verification. Single-cell foundation models trained on tens of millions of transcriptomes (scBERT [17], scGPT [18], Geneformer [19], scFoundation [20], UCE [21]) have raised the prospect of in silico screening at scale, and an ambitious programme—reviving the in-silico cell-modelling vision of the first whole-cell computational model [22]—now frames the *AI virtual cell* as a model that predicts cellular responses to unseen perturbations while flagging its own uncertainty [23]. Two obstacles stand between this prospect and phenotype verification. First, systematic benchmarking shows that state-of-the-art deep-learning models do not consistently outperform simple linear baselines at perturbation-effect prediction [24–26]; only recently has a model surpassed linear baselines at scale [27], and the most effective solutions inject structured domain knowledge as explicit features. Second, and more fundamental for the present problem, models such as scGen [28], CellOT [29], XPert [30] and PerturbNet [31] reconstruct transcriptome-wide responses but expose no phenotype-resolved interface, while transcriptome–language interfaces such as CellWhisperer [32], Cell2Sentence [33] and GenePT [34] and stepwise prediction frameworks such as VCWorld [35] render cell state language-compatible without closing the iterative loop these phenotypes require. None verifies whether a distributed, multicomponent phenotype—rather than the bulk transcriptome—actually moves under a candidate perturbation.

A parallel line of work couples predictive tools to large language model (LLM) agents that reason over the discovery process itself. ClockBase Agent [36] reanalyzed tens of thousands of intervention–control comparisons to surface age-modifying candidates; Google DeepMind’s Co-Scientist proposed and ranked hypotheses through a generate-debate-evolve architecture, recovering active compounds for acute myeloid leukemia [37]; FutureHouse’s Robin closed a literature-to-experiment loop over successive rounds to nominate ripasudil for dry age-related macular degeneration [38]; and Biomni [39] executed tasks autonomously across many biomedical domains. Related agents extend this paradigm to adjacent scientific tasks: tool-using chemistry agents such as Coscientist [40] and ChemCrow [41] plan and execute synthesis, and literature-synthesis agents such as PaperQA2 [42] retrieve and reason over the primary literature at expert level. These systems establish that LLM agents can traverse the hypothesis space of drug discovery productively. But they reason about candidates largely through qualitative literature association rather than quantitative prediction of the weak, module-level phenotype shifts that matter here, and—critically—they optimize against a fixed scoring objective that is never itself exposed to experimental test; even within the modelling literature, prevailing prioritization scores rely on heuristics that cannot be improved by experimental feedback [4]. The capability these phenotypes demand is reflexive quantitative reasoning: grounding symbolic hypotheses in numerical predictions of transcriptional outcome, and auditing whether the rule that defines a good candidate remains consistent with biology after each experimental round.

Here we present an AI-in-the-loop framework that makes phenotype verification explicit and the scoring objective revisable, integrating three components that each act as a verifier. PACE (Phenotype for Age-related Cell Evaluation) is a phenotype verifier: a panel of directionally scored, cell-type-resolved immune-aging gene sets selected for cross-cohort stability across four PBMC cohorts, which captures the distributed phenotype and outperforms established aging clocks in zero-shot benchmarking on in-house cohort of elderly donors. CellQ (Cell Quantization) is a virtual-cell verifier: a residual vector quantization (RVQ) tokenizer that compresses each single-cell transcriptome into eight discrete tokens aligned natively to the vocabulary of a reasoning LLM, with no cross-modal projection network, and predicts perturbation-induced module shifts autoregressively from few-shot prompts, verifying weak module-level responses where differential-expression recovery fails. An Analyzer–Planner–Auditor agent closes the loop at the level of the objective: after each experimental round it audits its own scoring assumptions against the wet-lab response matrix. Using primary human PBMCs from elderly donors as a prospective testbed, the agent screened 110 compounds, surfaced an asymmetric reversibility pattern that aged-down modules proved substantially more reversible than aged-up modules, and on that basis revised its scoring rule from an equal-weight average to a balance-constrained minimum, a refinement that generalized to an independent compound set and tracked functional T-cell aging readouts more closely than the original rule. The discovery objective thereby becomes an experimentally revisable hypothesis rather than a fixed target, defining a generalizable paradigm for phenotype-guided, module-resolved iterative discovery against complex transcriptional targets.

## Results

### Architecture of the verification-driven discovery agent

To address the verification problem posed by distributed, weakly-affected phenotypes, we designed an AI discovery agent in which the scoring rule that ranks drug candidates is itself revisable: each experimental round both prioritizes drugs and tests whether the current scoring rule is still consistent with biology (Fig. 1a). The architecture separates an outer hypothesis-management layer that may evolve between rounds from an inner workflow that we deliberately keep fixed. The outer hypothesis agent comprises three roles (Analyzer, Planner, and Auditor), which together read the round-*N* wet-lab response matrix, propose an updated scoring rule, and accept or reject the update against the evidence. At round 1 the same agent performs a cold-start from literature; we refer to this literature-derived starting point—a set of mechanistic hypothesis categories together with an initial equal-weight scoring rule—as Hypothesis 0. The Analyzer can additionally ingest drug analyses produced by external agentic systems such as OriGene [43].

**Fig. 1.**
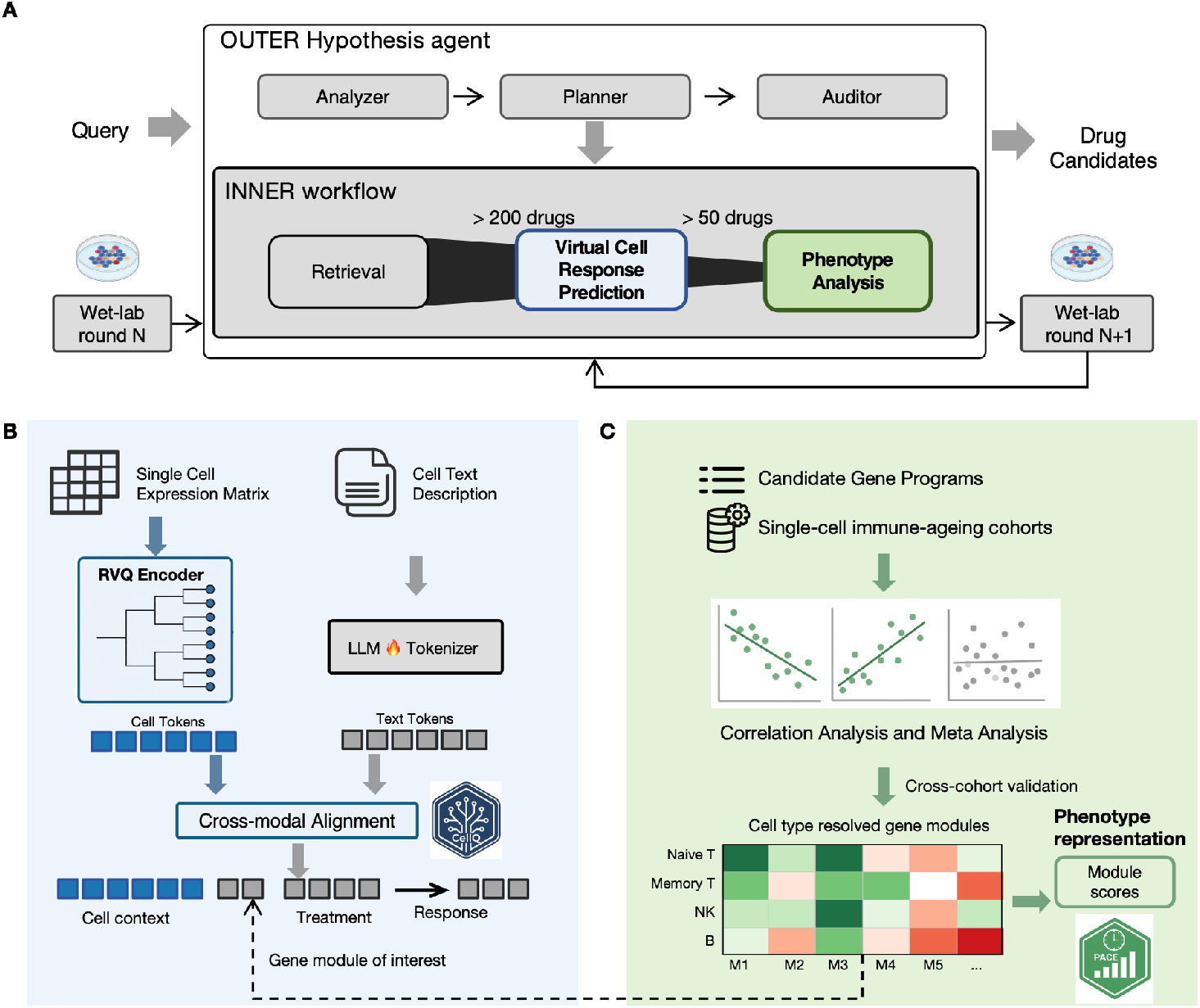
Overview of the AI-in-the-loop framework. **a**, Agentic architecture. The outer hypothesis agent (Analyzer, Planner, Auditor) reads the round-*N* wet-lab response matrix and updates the discovery objective for round *N* + 1, directing an inner workflow of retrieval, virtual-cell response prediction, and phenotype analysis. The framework is phenotype-agnostic; the light-blue and light-green modules are instantiated for immune aging by CellQ (**b**) and PACE (**c**). **b**, CellQ, a virtual-cell response engine instantiating the light-blue module in **a**. CellQ predicts perturbation-induced shifts at single-gene or module resolution given baseline cell context and a candidate compound. **c**, PACE, a phenotype engine instantiating the light-green module in **a**. PACE scores each candidate compound’s predicted response against a panel of cell-type-resolved, directionally scored immune-aging modules.

The inner workflow is engineered to be deterministic and replicable across rounds, with as few free variables as possible (Fig. 1a). It has three components. *Retrieval* pulls candidate drugs from a curated bank by cluster retrieval anchored on good-performing seed drugs (Methods). *Virtual-cell verification* is performed by CellQ, an RVQ-tokenized cell representation aligned to a reasoning LLM that is able to predict perturbation-induced shifts autoregressively from few-shot prompts (Fig. 1b, light blue); because the prediction unit is an LLM, the same prompt can ask for predictions at single-gene or module resolution, leveraging the LLM’s reasoning to combine context at either scale. *Phenotype verification* is performed by PACE, a panel of directionally scored, cell-type-resolved immune-aging modules against which the predicted module shifts are evaluated (Fig. 1c, light green). PACE supplies the scoring rule that the outer agent revises round-by-round and is the only phenotype-specific component; CellQ and the agent architecture apply unchanged to any phenotype expressible as a panel of directional, cell-type-resolved gene sets.

### RVQ–LLM enables cell representation and module-level perturbation verification

The virtual cell prediction model in the inner workflow was established first. The cell tokenizer uses residual vector quantization (RVQ) because the residual structure aligns naturally with the hierarchical organization of cell identity: each successive codebook refines a coarser context (Fig. 2a). The RVQ encoder was trained on more than 10^8^ single cells assembled from public atlases; eight residual codes proved sufficient to summarise a cell (Supplementary Fig. 1). After training, the code hierarchy exhibits an emergent organization across biological scales (Fig. 2b; Supplementary Fig. 2): CB0 captures broad cell context, CB1 cell identity, CB2 activation or resting state, CB3 functional programmes, and CB4 pathway-level signatures. This hierarchy was not pre-specified; it emerged spontaneously from the RVQ training objective.

**Fig. 2.**
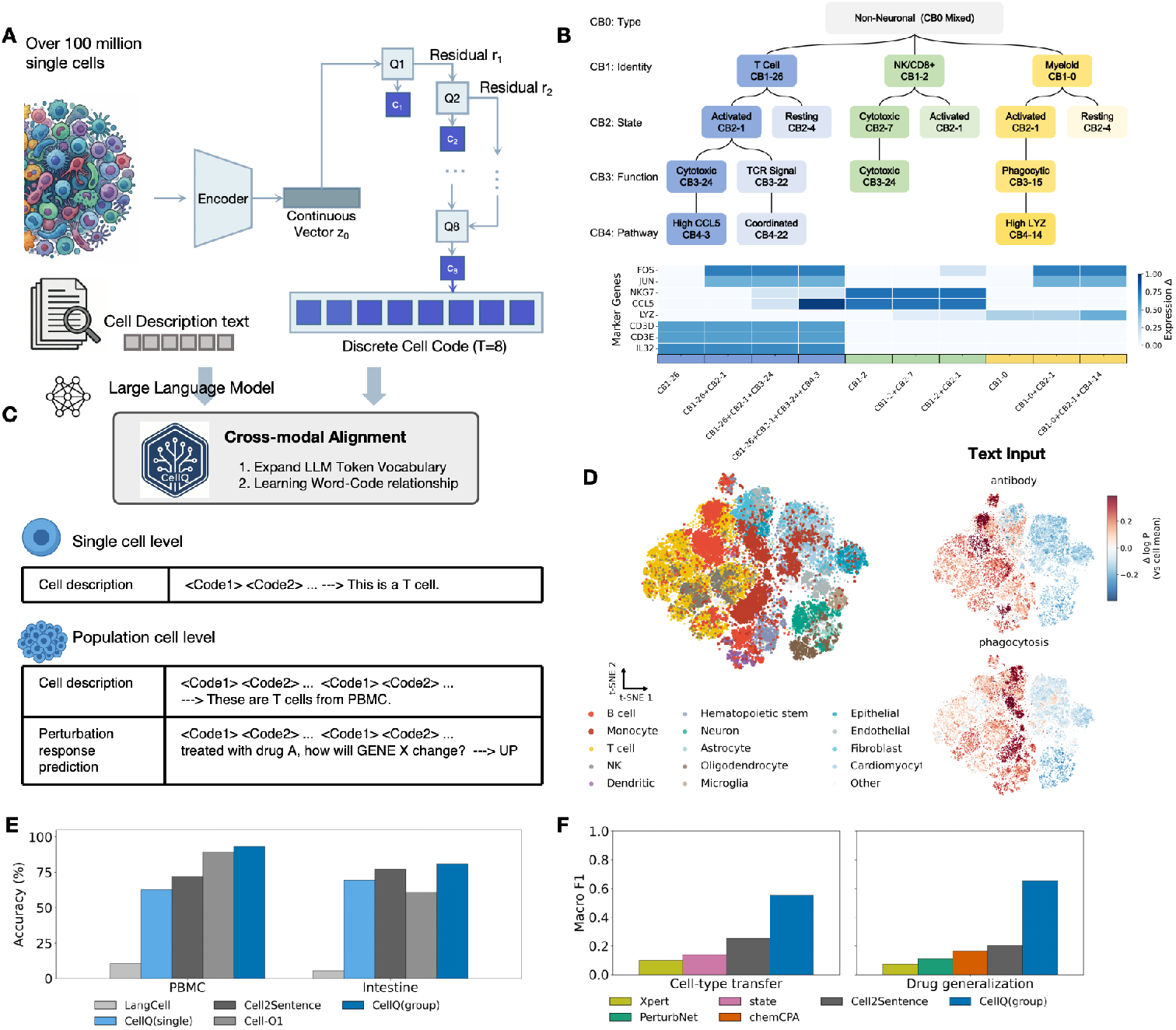
The RVQ–LLM framework and cross-modal alignment. **a**, Schematic of the RVQ– LLM pipeline. More than 10^8^ single cells are encoded into continuous latent vectors and quantized through residual vector quantization into discrete cell codes of depth *T* = 8. **b**, RVQ code-book organization across biological scales (CB0–CB4), with marker-gene heatmap for successive code combinations. **c**, Cross-modal alignment workflow. Code tokens expand the LLM vocabulary, and word–code relationships are learned through supervised fine-tuning to enable single-cell-level, population-level, and perturbation-level tasks. **d**, t-SNE visualization of cells from multiple datasets: (left) coloured by annotated cell type; (right) overlaid with query-conditioned text–cell alignment scores for “antibody” and “phagocytosis”. **e**, Cell-type classification accuracy on PBMC and intestine datasets. **f**, Perturbation benchmarking on the Tahoe dataset for cell-type-transfer and drug-generalization tasks.

A representative branch illustrates the emergent ordering at the gene level. CB1-26 marks T-cell identity, CB2-1 denotes an activated state, CB3-24 marks cytotoxic function and CB4-3 corresponds to a CCL5-high programme. The accompanying marker-gene heatmap shows that adding successive codes changes marker genes in a biologically coherent order, including T-cell markers *CD3D*, *CD3E* and *IL32*, immediate-early genes *FOS* and *JUN*, cytotoxic genes *NKG7* and *CCL5*, and myeloid-associated *LYZ* (Fig. 2b).

We then aligned the cell tokens with a reasoning-capable LLM. The RVQ codebooks were added directly to the LLM vocabulary, and word–code relationships were learned through supervised fine-tuning (Fig. 2c). The resulting system supports single-cell, population-level, and perturbation-level tasks through a unified chat template in which RVQ tokens and text tokens share the same embedding space, with no cross-modal projection network. Alignment quality was evaluated by t-SNE projection of cell embeddings, first coloured by annotated cell type to confirm that RVQ codes capture cellular identity, and then overlaid with query-conditioned scores for the text terms “antibody” and “phagocytosis” (Fig. 2d; Supplementary Fig. 3): antibody-related scores localize to B-cell-rich neighborhoods and phagocytosis-related scores to monocyte-rich neighborhoods, indicating that the alignment produces interpretable, biologically meaningful representations.

In cell-type classification, CellQ with group-code aggregation achieved the strongest performance on both PBMC and intestine benchmarks relative to LangCell, CellQ(single), Cell2Sentence, and Cell-01 (Fig. 2e; Supplementary Fig. 4). The compression advantage is a direct consequence of the RVQ design: each cell is represented by 8 tokens compared with 200+ tokens for gene-list-based methods such as Cell2Sentence. The resulting token efficiency allows more cells to be packed into the LLM context window, enabling richer population-level reasoning without token explosion, and motivates the use of group-level RVQ codes for downstream perturbation prediction.

For perturbation prediction, we used the Tahoe-100M single-cell perturbation atlas [44] with two task-specific test partitions: a *cell-type-transfer* split that holds out cell lines unseen during training, and a *drug-generalization* split that holds out compounds unseen during training. Predictions were obtained through few-shot prompting of the aligned RVQ–LLM with no perturbation-specific fine-tuning, leveraging the LLM’s in-context reasoning to map exemplar perturbations onto the queried compound. CellQ(group) achieved the highest macro-F1 in both settings, outperforming Xpert, PerturbNet, State, ChemCPA, and Cell2Sentence baselines (Fig. 2f; Supplementary Fig. 5). A non-RVQ LLM ablation that replaces RVQ codes with the bare LLM input performed substantially worse: RVQ discretization, not the LLM backbone alone, drives the gain.

### Directionally stable modules capture immune-senescence and enable hypothesis-driven screening

A robust phenotype representation is another prerequisite for verification. Many representations of immune aging have already been proposed, each derived from a different cohort or methodology — some from independent single-cohort analyses, others from merged multi-cohort sets — and each usually summarized as a gene set. These independently-derived gene sets overlap surprisingly little (Supplementary Figs. 6, 7), and no ground truth selects one of them as canonical. We therefore took a meta-analytic ensemble approach: assemble the gene sets these representations produced as candidates and identify the ones that are stable across cohorts.

PACE was constructed from three public discovery cohorts (Wang [45], AIDA [46], and OneK1K [47]; total *n* = 1,550 donors) with an independent in-house validation cohort of 434 elderly donors held out entirely for out-of-distribution (OOD) evaluation (Fig. 3a). The in-house atlas captures the major immune cell types (Fig. 3b). We collected 54 candidate gene sets from seven source categories spanning cell-type-specific and bulk-level methodologies (Fig. 3c) and filtered them by cross-cohort significance, direction consistency, and monotonicity with age (Supplementary Figs. 8, 9). Ten gene sets survived this filter and form the final PACE Aging Index. A forest plot of age coefficients (Fig. 3d) shows directional replication across all four cohorts: four modules decrease with age, while the other six increase with age across specific cell types. The composite PACE Aging Index increases monotonically across six chronological age bins in the OOD in-house cohort (Fig. 3e), and in zero-shot benchmarking against five established aging clocks achieves the highest Pearson correlation with chronological age (*r ≈* 0.40; Fig. 3f).

**Fig. 3.**
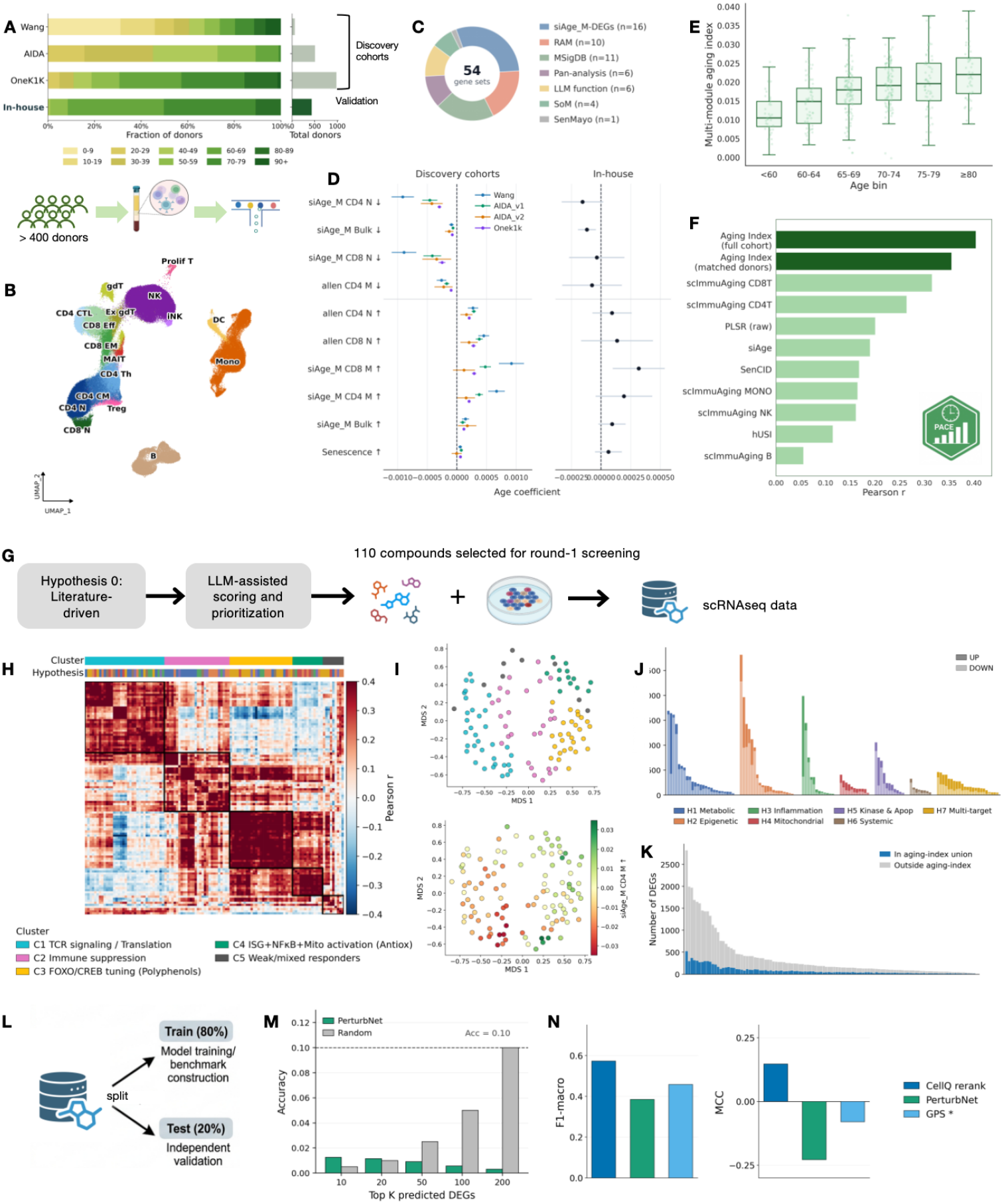
Construction and application of PACE, a multi-module aging index for immune-cell aging and drug screening. **a**, Age distribution of discovery cohorts (Wang, AIDA, and OneK1K; total *n* = 1,550) and out-of-distribution (OOD) in-house validation cohort (434 elderly donors), with schematic of PBMC isolation and scRNA-seq profiling. **b**, UMAP of in-house OOD validation scRNA-seq data annotated by immune cell type. **c**, Composition of 54 curated gene sets by source category used for PACE construction. **d**, Forest plot of linear-regression age coefficients for ten representative candidate gene sets across discovery cohorts and OOD in-house validation (black dots). **e**, Multi-module PACE Aging Index versus chronological age across age bins in the OOD in-house cohort. **f**, Zero-shot benchmarking of aging clocks on held-out in-house data. **g**, Round 1 compound-selection workflow: literature-derived Hypothesis 0, LLM-assisted scoring and prioritization, and scRNA-seq profiling of 110 selected compounds. **h**, Pearson correlation heatmap of drug transcriptional responses clustered into five functional response groups (C1–C5), with an H1–H7 hypothesis-category annotation strip. **i**, Two stacked MDS scatter plots over the same compound space: (top) drugs coloured by transcriptional response cluster (C1–C5); (bottom) coloured by the *siAge M CD4 M ↑* aging-reversal score. **j**, Bar plot: number of up- and down-regulated DEGs per compound, coloured by hypothesis category. **k**, Overlap between compound-induced DEGs and the PACE aging-index gene union. **l**, Train–test data split schematic (80/20) for model-training benchmark construction and independent validation. **m**, Top-*K* predicted-DEG accuracy of PerturbNet versus random baseline. **n**, Module-level prediction performance of CellQ re-rank, PerturbNet, and GPS*, measured by MCC and ranking metrics.

With PACE and CellQ in place, we generated the first round of wet-lab data. The *≈*5,000-compound candidate pool was narrowed to a 110-compound panel through LLM-assisted scoring against the seven mechanistic categories of Hypothesis 0 (Fig. 3g): H1 Metabolic, H2 Epigenetic, H3 Inflammation, H4 Mitochondrial, H5 Kinase and Apoptosis, H6 Systemic, and H7 Multi-target (Supplementary Fig. 10).

Correlation analysis organizes the 110-compound response atlas into five transcriptional response clusters: C1 TCR signaling / translation, C2 immune suppression, C3 FOXO/CREB tuning (polyphenols), C4 ISG+NF-*κ*B+mitochondrial activation (antioxidants), and C5 weak or mixed responders, with an overlaid annotation strip showing the H1–H7 hypothesis category of each drug (Fig. 3h; Supplementary Fig. 11). Two stacked MDS scatter plots over the same compound space confirm that this cluster structure is functionally meaningful: drugs separate cleanly by transcriptional response cluster (top, coloured by C1–C5), and the same MDS is organized by the *siAge M CD4 M ↑* aging-reversal score (bottom; Fig. 3i; Supplementary Figs. 12, 13).

Across the panel, differentially expressed gene (DEG) counts vary substantially by compound and category, with H1, H2, and H3 compounds showing the most prominent transcriptional response ranges (Fig. 3j). However, only a minority of drug-induced DEGs overlap the PACE aging-index gene union; most DEGs fall outside the aging-index set entirely (Fig. 3k). This implies that single-gene-level prediction is unlikely to recover the relevant pharmacological signal, and motivates evaluating CellQ at the module level rather than the DEG level.

To test whether CellQ’s predictions are biologically meaningful, we benchmarked CellQ re-rank against three transcriptome-guided methods (PerturbNet, GPS*^∗^*, and DrugReflector) on a held-out 22-drug subset of the 110-compound panel (Fig. 3l). PerturbNet was trained on the remaining 88 drugs; GPS*^∗^* and DrugReflector are zero-shot, comparing each drug’s reference transcriptional signature against a young-vs-old immune signature derived from PACE, and never seeing the drug-treatment data. All methods were scored on the same per-(drug, module) BETTER/WORSE task across the 22 held-out drugs and 10 PACE modules. PerturbNet’s top-*K* predicted-DEG accuracy on the held-out fold was at or below the random baseline across *K* = 10– 200 (Fig. 3m; Supplementary Fig. 14): raw DEG recovery is not a sufficient surrogate for module-level prioritization. At the module level, CellQ re-rank achieves the best macro-F1, and remains the best under the stricter Matthews correlation coefficient (MCC; Fig. 3n; Supplementary Fig. 15).

### Mismatch-driven iterative discovery

With PACE and CellQ in place, we tested whether the full inner workflow enriches for better-performing drugs and whether the outer agent revises the scoring rule when the data demand. Under leave-one-out evaluation on the 110-drug round-1 panel (Fig. 4a), cluster retrieval achieved significantly higher Normalized Discounted Cumulative Gain (NDCG) than random retrieval at both NDCG@5 and NDCG@10 (Fig. 4b). The CellQ re-rank that converts each retrieval shortlist into per-(drug, module) BETTER/WORSE calls is well-calibrated, with precision rising monotonically with the model’s self-assessed confidence for both classes (Fig. 4c). Ranking the round-1 candidates by predicted retrieval score and computing signed enrichment per hypothesis category, we recovered the known mechanistic structure even though retrieval never sees mechanism labels: mitochondrial protectors (H4) concentrated at the top of the ranking, and metabolic regulators (H1) at the bottom, with weaker positive enrichment for epigenetic (H2) and inflammation (H3) compounds (Fig. 4d).

**Fig. 4.**
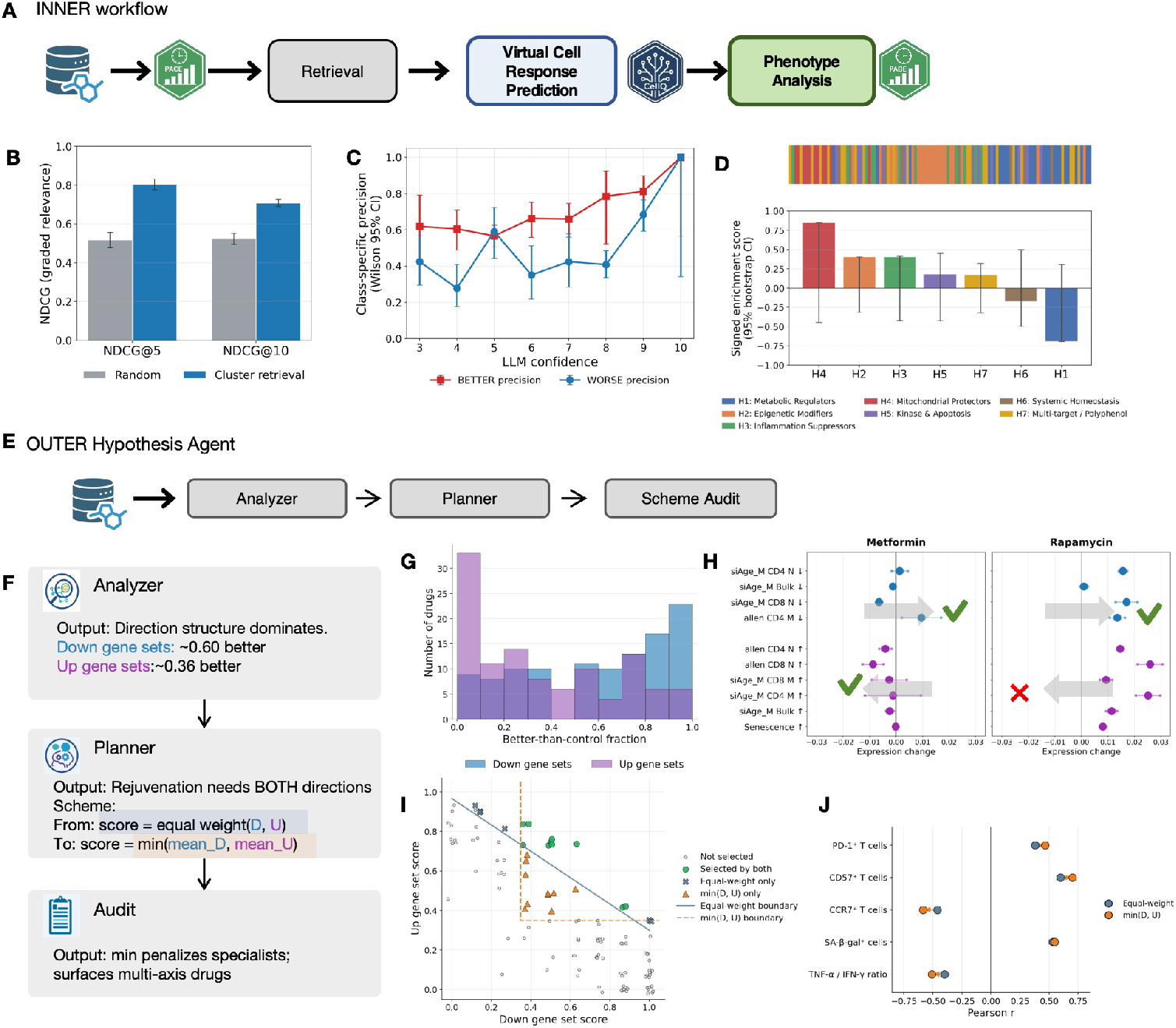
Validation of the inner retrieval–prediction workflow and the outer hypothesis self-audit. **a**, Inner-workflow schematic: retrieval, CellQ virtual-cell response prediction, phenotype scoring. **b**, NDCG@K under leave-one-out on the 110 round-1 compounds. Error bars are paired-bootstrap 95% confidence intervals. **c**, CellQ re-rank calibration on 110 *×* 10 = 1,100 (drug, module) calls: precision of BETTER and WORSE calls versus self-assessed confidence, with Wilson 95% confidence intervals; the dashed line marks the empirical BETTER base rate. **d**, Signed enrichment of mechanistic categories (H1–H7) along the retrieval-score ranking of round-1 candidates, with drug-level bootstrap 95% confidence intervals. **e**, Outer hypothesis agent: Analyzer, Planner, Auditor. **f**, Agent output example: from Analyzer, to Planner, to Auditor. **g**, Distribution of restored aged-down and aged-up modules across the compound panel. **h**, Direction-resolved response profiles for metformin and rapamycin across PACE modules. **i**, Scatter plot of drug performance by aged-up and aged-down modules, and grouped by equal weight or MIN-combiner. **j**, Independent T-cell functional validation of 13 compounds: Spearman correlation of the equal-weight and MIN-combiner scores with PD-1^+^, CD57^+^, and CCR7^+^ T-cell fractions, SA-*β*-gal^+^ cells, and the TNF-*α*/IFN-*γ* ratio compared to the control.

The outer hypothesis agent then ingested the round-1 response matrix and worked through the Analyzer–Planner–Auditor cycle (Fig. 4e). The Analyzer surfaced a structural mismatch between the equal-weight scoring rule and the data: aged-down modules were substantially more reversible than aged-up modules (Fig. 4f), an asymmetry that held across the full compound panel (Fig. 4g) and at single-compound resolution, with metformin restoring both direction classes as expected while rapamycin only restores aged-down modules (Fig. 4h). Because immunorejuvenation should restore both direction classes simultaneously, an equal-weight additive rule was direction-blind: it ranked a direction-specialist on par with a balanced multi-axis drug.

The Planner therefore proposed a balance-constrained replacement (Hypothesis 1), min(*D, U*), which penalises direction specialists and surfaces multi-axis candidates (Fig. 4i), and the Auditor approved the revision. We locked the MIN rule and tested it against the equal-weight rule on an independent T-cell functional assay of 13 compounds (Methods): the MIN rule produced stronger and more consistent associations with the functional readouts (Fig. 4j; Supplementary Fig. 16). The scoring rule is therefore itself a falsifiable experimental object — the agent revises it when the response matrix demands, and the discovery target evolves round-by-round under the same architecture.

## Discussion

This work shows that an AI system can treat its own discovery objective as a testable claim rather than a fixed target. In conventional drug-repurposing pipelines, the scoring function is specified once by a human investigator and held constant throughout the search. Here, the Auditor examined the empirical reversibility pattern and identified a structural inconsistency between the initial additive scoring and the biological goal of multi-axis rejuvenation. The Planner then replaced equal weighting with a balance-constrained MIN rule. In effect, the discovery objective became an experimental object: a hypothesis about what immune rejuvenation should look like was exposed to cellular data, found wanting, and revised. This is the central methodological move of the work.

One biological finding emerged from this self-audit: aged-down modules are substantially more reversible than aged-up modules. Current aging clocks and rejuvenation scores typically treat all age-associated changes as equally targetable, but our data suggest that the transcriptional programmes of immune aging are not symmetric. Some axes resist pharmacological reversal more than others. That aspects of cellular aging are reversible at all is consistent with rejuvenation strategies such as partial and transient reprogramming, which restore youthful features without erasing cell identity [48, 49], and senolytic clearance of senescent cells [50]; our data add that, within a single phenotype, reversibility is itself direction-dependent. The MIN combiner penalizes compounds that restore only the more reversible modules, and rewards those that restore both directions. This asymmetry was not obvious from the literature or from the initial hypothesis design; it became visible only when the Auditor examined the experimental response matrix and detected the mismatch between the scoring function and the biological pattern.

For AI-driven science, this work demonstrates a capability beyond hypothesis ranking: reflexive quantitative reasoning. Here the agent surfaced a quantitative mismatch between the equal-weight scoring rule and the asymmetric reversibility pattern in the round-1 data, and replaced the rule with a balance-constrained alternative. This is a step toward AI systems that can surface their own quantitative inconsistencies, though its critiques remain bounded by the expressiveness of the scoring functions it can formulate.

This capability connects to a broader research direction in artificial intelligence: systems that can recursively examine and improve their own reasoning. Self-improvement can be framed as a Generator–Verifier–Updater (GVU) cycle, in which an agent generates proposals, an independent verifier evaluates them, and an updater revises the agent’s policy. The pattern has been demonstrated in mathematical discovery, where systems such as AlphaEvolve autonomously propose, test, and refine algorithmic solutions to open problems [51], and in language models that critique and revise their own outputs—through self-refinement [52], constitutional self-critique [53] and self-rewarding training [54]—or that automate the research cycle end to end, as in The AI Scientist [55]. The Analyzer–Planner–Auditor architecture described here instantiates one cycle of this pattern in a more demanding domain: verification is a biological measurement, the transcriptional response of primary human cells to pharmacological perturbation, rather than a computational metric. The Planner generates scoring rules (Generator), the Auditor evaluates them against experimental data (Verifier), and the revised objective is deployed in subsequent rounds (Updater). Unlike self-improvement systems that modify their own source code, the agent operates at the level of its discovery objectives, the scoring functions that encode what it means for a drug to be effective. This restriction is deliberate: confining self-modification to the objective layer, and grounding verification in experimental data, keeps the system interpretable and auditable by human scientists while still closing a self-improvement loop that current AI scientists cannot.

The broader significance of this work lies at the intersection of three converging research fronts. GPS and DrugReflector established that transcriptomic objectives can guide chemical search, but both assume that the desired transcriptional outcome is known in advance, and signature-reversion scoring can moreover reward generic anti-proliferative effects rather than target-specific reversal [56]. Co-Scientist and Robin demonstrated that LLM agents can productively traverse hypothesis spaces, but without quantitative perturbation models, they cannot verify whether a candidate drug would actually produce the desired transcriptional state. And the recursive self-improvement paradigm in AI—exemplified by Generator–Verifier– Updater cycles—has largely been demonstrated in software domains where verification is code execution. Our framework integrates these fronts: it provides transcriptomic objectives that are revisable (PACE), quantitative perturbation verification (CellQ), and objective-level self-audit grounded in experimental truth (Auditor), grounding a self-improvement loop in biological measurement rather than code execution, constrained not by compilers but by the transcriptional response of primary human cells. The clinical maturity of AI-driven drug discovery is also advancing in parallel [57], providing clinical validation that AI-driven discovery can produce candidates that succeed in human trials.

Several limitations warrant discussion. RVQ compression to eight discrete tokens necessarily discards fine-grained expression information, and systematic characterization of this information-loss boundary remains an open task. Our validation focused on a single phenotype, immune aging in PBMCs, chosen for its stringency as a multi-component, cell-type-resolved target. The framework’s modular architecture accepts any phenotype definition specified as gene sets, but empirical demonstration across disease areas awaits future work. The wet-lab scale, though sufficient for closed-loop self-correction, is modest relative to deployable chemical space, and the current demonstration ran a single revision round.

Beyond immune aging, the framework is designed to generalize: CellQ was pretrained on more than 10^8^ cells spanning diverse tissues and disease contexts, so its discrete token representation captures transcriptional states across lineages, and only the verification module (PACE here) is phenotype-specific and needs to be re-specified for a new target. Whether comparable directional asymmetries arise in other phenotypes such as fibrosis, neurodegeneration, or metabolic disease is an open question the same self-audit could address. More broadly, the same architecture is not specific to drug discovery: wherever AI systems make quantitative recommendations—in materials design, clinical-trial optimization, or environmental modeling—it could let agents examine whether their evaluation criteria remain consistent with observed outcomes, moving the recommendation objective from a fixed target to an empirically revisable hypothesis. Realizing this will require iterating the audit across multiple rounds, automated wet-lab execution, broader pharmacological coverage in perturbation prediction, and expansion of the audit scope from scoring-function critique to experimental-design critique. As LLM agents become more capable of traversing hypothesis spaces, the bottleneck shifts toward verification—the quantitative tools with which an agent can test its own assumptions against experiment—which is what this architecture supplies; as those models become increasingly powerful, such a framework will help close the gap toward fully autonomous scientific discovery.

## Methods

### RVQ–LLM virtual-cell model

#### Pre-training data

The cellular-state encoder was pre-trained on a single-cell transcriptomic compendium of more than 10^8^ human cells assembled from public atlases. The data span 105 dataset shards drawn from diverse tissues, age groups, and healthy or disease contexts. Counts were normalised to a common gene panel and log1p-transformed. Train and held-out splits were stratified at the dataset level so that no dataset appeared in both splits.

#### Cellular state encoding via residual vector quantization

A residual vector quantization (RVQ) autoencoder maps each cell’s gene-expression vector **x** *∈* R*^G^*to a latent state **z** *∈* R*^d^*through a multilayer-perceptron encoder, and approximates **z** as the sum of *L* = 8 codes drawn from independent codebooks of size *K* = 32,

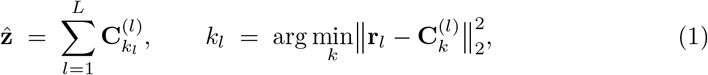

with residuals **r**_1_ = **z** and 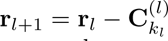 Each cell is summarised by an ordered *L*-tuple (*k*_1_*, …, k_L_*) that is consumed as a sequence of discrete tokens by the downstream language model. Lower codebooks capture coarse cell-identity axes (lineage, tissue compartment) while higher codebooks resolve fine-grained activation, cell-cycle, and metabolic states. A symmetric MLP decoder reconstructs gene expression from **z**̂.

The encoder is trained as a denoising autoencoder under a composite objective

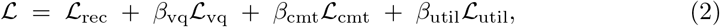

where 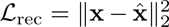 is the reconstruction loss, 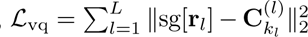 updates the codebook entries via the straight-through estimator, 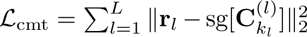 encourages the encoder to commit to the quantized representation, and *L*_util_ is a codebook-utilisation term that penalises collapsed codebook entries and promotes uniform usage across all *K* codewords at each level.

#### Language-model alignment

After RVQ pre-training, we aligned the LLM (Qwen3-4B-Thinking-2507) to the discrete cell-state space following the RVQ-Alpha [58] protocol, which combines continued pretraining with supervised fine-tuning. The eight residual codebooks are added directly to the LLM vocabulary in fixed positions; cell tokens and natural-language tokens therefore share a single embedding table and a single autoregressive objective, with no cross-modal projection network. Supervised fine-tuning followed an Evidence-First Feature-Injection strategy. Each training prompt presents the RVQ token tuple (*k*_1_*, …, k_L_*) together with the cell’s top-*K* most strongly expressed genes, and the target completion is a chain-of-thought that lists gene-level evidence before stating the conclusion (cell type, tissue, disease context, age group, characteristic markers). This evidence-before-conclusion structure binds the newly added RVQ tokens to gene-level semantics through the language-modelling loss itself, in place of a contrastive or projection-based aligner.

#### Perturbation prediction via few-shot prompting

Compound-induced transcriptional shifts were predicted through few-shot prompting of the aligned RVQ–LLM model, with no perturbation-specific fine-tuning. The setup follows the VCWorld few-shot paradigm [35], augmented with discrete cell tokens: the baseline cell population is summarised by its group-level RVQ codes and inserted into the prompt alongside a small set of perturbation exemplars retrieved from the training split, and the model autoregressively predicts the post-perturbation differential expression. Generalisation to unseen perturbations is achieved through retrieval and in-context exemplars; no parameter updates are made between queries.

The key evaluation metric is not whole-transcriptome reconstruction error but module-level verification. Formally, for a compound *c* and a PACE module *m*, let the observed module shift be 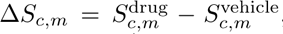, where *S_c,m_* denotes the UCell module score. For an aged-up module (positive age coefficient *d_m_* = +1), a beneficial shift corresponds to *−*Δ*S_c,m_* (reversal of the aging direction); for an aged-down module (*d_m_* = *−*1), a beneficial shift corresponds to +Δ*S_c,m_*. The binary classification target is 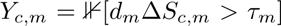, where *τ_m_* is a donor-stratified threshold derived from the vehicle-control variance. CellQ’s task is to predict *Y_c,m_*for each (compound, module) pair. DEG exemplars in the prompt provide coarse perturbation context (the compound’s known transcriptional fingerprint), but evaluation is performed on the full module including genes that are not among the top DEGs for that compound. This design allows CellQ to leverage the LLM’s flexible module-level reasoning to predict coordinated shifts beyond DE gene recovery.

#### Benchmarking

We compared the full RVQ–LLM model against ablations and external baselines on two task families. For cell-type classification, CellQ was evaluated against LangCell, CellQ(single), Cell2Sentence, and Cell-01 on PBMC and intestine reference datasets (Fig. 2e). For perturbation prediction we used the Tahoe-100M single-cell perturbation atlas with a shared training partition and two task-specific test partitions: a *cell-type-transfer* split that holds out cell lines unseen during training, and a *drug-generalization* split that holds out compounds unseen during training. CellQ was benchmarked against Xpert, PerturbNet, State, ChemCPA, and Cell2Sentence under each split (Fig. 2f). A non-RVQ ablation that replaces RVQ codes with the bare LLM input was included to isolate the contribution of discrete tokenisation.

### Phenotype abstraction and directional scoring

#### Data collection and processing

We defined computable immune-aging phenotype axes from four independent PBMC single-cell transcriptomic cohorts with donor age metadata: a repertoire-integrated cohort with scRNA-seq, T/B cell receptor sequencing and mass cytometry (Wang) [45]; the Asian Immune Diversity Atlas (AIDA) [46]; an immune eQTL cohort mapping autoimmune disease genetics (OneK1K) [47]; and a local elderly cohort generated in-house. Because the Asian immune diversity cohort was represented by two data releases of the same underlying sample collection, we treated these as a single discovery stratum in meta-analysis to avoid sample overlap.

For each cohort, we computed gene-set activity using UCell [59], a rank-based single-cell enrichment score, and aggregated to donor-level summaries. Among single-sample gene-set scoring methods descending from gene-set enrichment analysis [60]— including GSVA [61] and AUCell [62]—we selected the rank-based UCell score for its robustness to the sparsity and dropout characteristic of single-cell data, consistent with benchmarks of single-cell gene-set scoring [63]. Within each cohort, we fit gene-set-specific linear models with donor-level score as the response and chronological age as the predictor, adjusting for sequencing depth and batch where batch structure was informative. We extracted the age coefficient, standard error, and *p* value per gene set per cohort, and combined cohort-level effects by random-effects meta-analysis with Benjamini–Hochberg false-discovery-rate control. Candidate gene sets were then admitted under one of two categories: *Core*, requiring per-cohort partial-conjunction *m* = 4*/*4 (significant in the expected direction in all four discovery datasets) with median *|t| ≥* 2.0 and Spearman age–score monotonicity in at least three of four datasets; and *Extension*, requiring partial-conjunction *m* = 3*/*4 with the same monotonicity threshold, manually finalised so that the panel covers both naive and memory states for both CD4 and CD8 T cells. The resulting axes were annotated as aged-up or aged-down based on the meta-analytic effect sign.

#### Age clocks benchmarking

We benchmarked these axes against established aging and senescence scores (PBMC cell-type proportion clock [64], sc-ImmuAging [10], siAge [45], hUSI [11], and SenCID [12]) to ensure they captured biologically relevant aging signal rather than technical noise. Pearson correlations with chronological age were computed on the full in-house validation cohort (PACE Aging Index) and on a matched-donor subset restricted to donors with every benchmarked clock defined (matched-donor PACE variant); results are reported in Fig. 3f.

#### Drug-treated scRNA-seq data analysis

For each (donor, compound), we counted the ten PACE modules on which the compound was classified BETTER, where BETTER requires the donor-stratified module score to shift in the anti-aging direction by more than the vehicle-control threshold. The per-compound donor-aggregated restoration rate *u̅* is the mean across donors of the fraction of restored modules. Compounds were split at the lower and upper quartiles of *u̅* across the 110-drug panel into tiers T1 (top 25%), T2 (interquartile 50%) and T3 (bottom 25%).

#### Transcriptome-guided methods comparison

All wet-lab comparisons in this section use the per-(drug, PACE axis) BETTER/WORSE labels defined in §7. We compared CellQ rerank against three published transcriptome-guided methods that operate at the gene level: GPS [3], which scores compounds by transcriptomic reversal against a target signature (*Z*_RGES_); DrugReflector [4], which runs active learning over a pre-computed L1000 perturbation library [65]; and PerturbNet [31], a deep-learning model trained to predict the post-perturbation transcriptome.

CellQ rerank predicts a BETTER or WORSE outcome for each (compound, PACE module) pair by few-shot prompting [35], with four task-specific components inserted into the prompt: the query drug’s targets and pathways from PrimeKG [66]; an experimental-context cue restricting reasoning to cell-autonomous mechanisms in an in vitro PBMC culture; the target module’s representative genes and its aging direction; and reference compounds retrieved by multi-modal similarity from the round-1 response matrix, each annotated with its observed BETTER or WORSE label. The retrieved exemplar set is balanced so that both outcome classes are represented. The model returns a decision, a 1–10 confidence, and a one-sentence rationale.

Each method was applied to the round-1 compound panel under matched conditions. PerturbNet was trained on 88 of the 110 compounds and held out on the remaining 22 (Fig. 3l). GPS*^∗^* and DrugReflector are zero-shot: they compare each drug’s reference signature against a young-vs-old immune signature derived from PACE without ever seeing the drug-treatment data. For both zero-shot methods, compound-level scores (GPS*^∗^*: *Z*_RGES_; DrugReflector: L1000 library similarity) were converted to per-(drug, module) BETTER/WORSE calls by a per-module median split. DrugReflector’s pre-computed L1000 library covers 10 of the 22 held-out drugs; the remaining 12 lack a prediction (coverage gap). Predictions were evaluated on the 22 held-out drugs across the 10 PACE modules using macro-F1, MCC, and Cohen’s *κ* at the BETTER/WORSE level, and against per-gene differential expression using top-*K* accuracy at *K* = 10–200. The same evaluation protocol was applied to CellQ re-rank.

The Matthews correlation coefficient is defined as

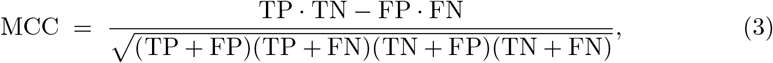

where TP, FP, TN, FN are the counts in the confusion matrix at the (drug, module) level. MCC ranges from *−*1 (perfect anti-correlation) to +1 (perfect agreement), with 0 at chance. Unlike macro-F1 — which averages per-class F1 and remains positive whenever any class is predicted at all — MCC penalises both false positives and false negatives jointly and turns negative when a predictor is systematically miscalibrated against the truth (for example, predicting BETTER on drugs that are wet-lab WORSE at a rate above the per-module base rate). This is the failure mode that distinguishes a method whose macro-F1 looks acceptable but whose decisions are anti-correlated with the wet-lab ground truth.

### Agentic screening and hypothesis update

#### Round-1 candidate retrieval and prioritization

The round-1 candidate pool was assembled from two public sources: the Target-MOI L9200 small-molecule library and DrugBank [67], totalling approximately 5,000 compounds. Candidates were filtered for tractability and prior safety flags, then prioritized by LLM-assisted scoring against the seven mechanistic categories of the literature-derived Hypothesis 0 (H1 Metabolic, H2 Epigenetic, H3 Inflammation, H4 Mitochondrial, H5 Kinase and Apoptosis, H6 Systemic, H7 Multi-target). The 110-compound panel taken to wet-lab screening (Fig. 3g) was selected to balance hypothesis coverage and prior literature evidence of immune relevance.

#### Inner screening agent

The inner layer is a two-stage retrieval–prediction pipeline evaluated by leave-one-out (LOO) on 110 wet-lab-tested anti-aging compounds. Each drug is first rendered into a short hybrid mechanism prompt of the form “*¡Name¿ is a ¡category¿ drug with ¡MoA-category¿ mechanism. ¡first sentence of the curated mechanism description¿.*” Prompts are wrapped with the Qwen3-Embedding retrieval instruction template and encoded with Qwen3-Embedding-4B, a 4B-parameter decoder-only encoder pretrained with a contrastive retrieval objective. We apply last-token pooling on the final hidden state with left-padding (the canonical configuration distributed with the model), followed by L2 normalization, yielding 2,560-dimensional unit vectors. Under LOO, the 109 held-in drugs are partitioned by K-Means into ten prototype clusters; centroids are re-normalized to unit norm, and each cluster is summarized by the mean restoration rate *u̅* (defined in §7) of its members. For each held-out query drug, the predicted *u̅* is the cluster mean of its nearest centroid by cosine similarity. The shortlisted candidates are then re-ranked by CellQ: for each (drug, module) pair the bank is queried for the top-6 mechanistically similar exemplars under a balanced strategy (at least two minority-class exemplars), and the model is prompted to return a forced BETTER/WORSE decision with a self-assessed confidence on a 1–10 scale. Per-drug aggregate scores are obtained by combining decisions across the ten modules with the scoring rule produced by the outer hypothesis agent (equal-weight mean in round 1, MIN combiner after the round-1 self-audit).

#### Inner-workflow evaluation

Retrieval was evaluated under LOO on the 110-drug benchmark using NDCG@K (Normalized Discounted Cumulative Gain), with graded relevance T1=2, T2=1, T3=0 from the donor-aware drug response tiers. Significance was assessed by paired bootstrap (*n* = 2,000) resampling the 110 drugs and computing the matched difference NDCG@K(retrieval) – NDCG@K(random) within each bootstrap sample. The CellQ re-rank in Fig. 4c was evaluated on the same 110-drug LOO pool (110 *×* 10 = 1,100 (drug, module) cells); per-confidence-bin precision was computed separately for BETTER and WORSE calls with Wilson 95% confidence intervals.

#### Mechanism-structure recovery

The round-1 candidate compounds were ranked by their predicted retrieval score, and per-hypothesis-category signed enrichment statistics (H1–H7) were computed by running-sum gene-set-enrichment-style scoring over this ranking. Drug-level bootstrap (*n* = 1,000) provided 95% confidence intervals on the enrichment score, and label-permutation testing (*n* = 2,000) provided category-level *p* values.

#### Outer hypothesis agent

After each experimental round, the outer layer analyses the response matrix to update the discovery objective. It comprises three roles, each implemented by prompting the underlying LLM with a role-specific definition of title, expertise, goal, and input/output. All agent outputs, prompt versions and tool calls were logged for audit and are available with the code release.

#### Analyzer

Reads the round-*N* wet-lab response matrix and compares responses across drugs, donors, doses, and phenotype directions to identify reproducible response archetypes and any mismatch between the current scoring rule and the data. The Analyzer can run horizontal analyses across compounds and vertical analyses that link differentially expressed genes to pathways and literature mechanisms for individual drugs.

#### Planner

Proposes an updated scoring rule and a next-round retrieval query based on the mismatches surfaced by the Analyzer.

#### Auditor

The Auditor checks whether the Planner’s proposal is grounded in evidence the Analyzer actually surfaced — that each substantive element of the proposed scheme cites the specific findings used to justify it, and that those findings plausibly support what they back. Proposals lacking this grounding are returned to the Planner for revision.

#### Hypothesis-update rule

The initial scoring objective used an equal-weight average of aged-down-axis restoration and aged-up-axis reversal:

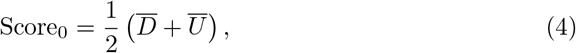

where *Dᐨ* is the mean improvement score for aged-down modules and *Uᐨ* is the mean improvement score for aged-up modules. Round 1 wet-lab validation revealed structured asymmetric reversibility: aged-down axes were more frequently improved than aged-up axes were reversed. The Planner therefore proposed, and the Auditor retained, a balance-constrained objective:

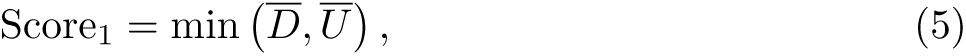

subject to viability and immune-function filters. This rule penalizes single-direction specialists and prioritizes candidates that improve both direction classes. The updated objective was then used to reprioritize candidates for the next experimental round without retraining the underlying RVQ–LLM model.

### Wet-lab perturbation experiments

#### PBMC source and culture

Cryopreserved peripheral blood mononuclear cells (PBMCs) were obtained from STEMCELL Technologies (samples collected under approved ethical protocols with documented informed consent). PBMCs were thawed, recovered briefly in pre-warmed medium, and seeded into 96-well ultra-low-attachment plates at 4 *×* 10^5^ cells per well in RPMI 1640 (Gibco, 61870-036) supplemented with 10% fetal bovine serum (FBS; VivaCell, C04002) and penicillin/streptomycin (Gibco, 15140-122). After 48 hours the medium was refreshed and compounds were added at the final concentrations indicated for each experiment, with half-volume medium replacements daily. Cultures were maintained for 4 days before harvest. This protocol was shared between the round-1 transcriptomic screen and the targeted T-cell rejuvenation assay; the two experiments differ in their compound panel and downstream readout.

#### Round-1 transcriptomic screen (110 compounds)

Three donors were used for the round-1 screen. PBMCs were cultured and treated as described above with the 110-compound panel at the final concentrations listed in Supplementary Table 1. After 4 days of treatment, cells were harvested and processed for single-cell RNA sequencing using 10x Genomics 3’ chemistry with Cell Multiplexing Oligo (CMO) tagging for sample multiplexing and donor demultiplexing in Cell Ranger. Reads were aligned to GRCh38 and quantified with Cell Ranger.

#### T-cell rejuvenation assay

For an independent functional validation, 13 compounds were selected and profiled in a T-cell-specific assay (Supplementary Table 2). Primary T cells were purified from cultured PBMCs using the Human CD3^+^ T Cell Isolation Kit (Yeasen, 37651ES60) and expanded in ImmunoCult-XF T Cell Expansion Medium (STEMCELL Technologies, 10981) supplemented with 10 ng/mL recombinant human IL-2 (PeproTech, 200-02-10UG) and stimulated with ImmunoCult Human CD3/CD28 T Cell Activator (STEMCELL Technologies, 10991). After 7 days of expansion, T cells were re-seeded into 96-well ultra-low-attachment plates at 4.5 *×* 10^5^ cells per well in RPMI 1640 with 10% FBS, 5 ng/mL IL-2 and penicillin/streptomycin, and treated with the 13 compounds for 4 days. T cells were then harvested for phenotypic and senescence profiling, and the matched culture supernatants were collected for cytokine ELISA. For phenotyping, 2 *×* 10^5^ T cells per condition were washed in 1% BSA / DPBS and stained on ice for 30 min with a fluorochrome panel: PE-Cy7 anti-human CD3 (BD Biosciences, 563423, 1:200), PE anti-human CCR7/CD197 (BioLegend, 353203, 1:200), APC anti-human PD-1/CD279 (BioLegend, 621609, 1:200), Brilliant Violet 711^TM^ anti-human CD57 (BioLegend, 393328, 1:200), and the LIVE/DEAD^TM^ Fixable Near-IR (780) Viability Kit (Invitrogen, L34992) for dead-cell exclusion. Data were acquired on a CytoFLEX LX flow cytometer (Beckman Coulter) and analysed in FlowJo v10.8.1; the proportions of PD-1^+^ and CD57^+^ cells within the viable T-cell compartment were quantified as measures of exhaustion and terminal differentiation, hallmarks of T-cell immunosenescence [68] (gating strategy in Supplementary Fig. 17). Parallel aliquots were stained with the CellEvent^TM^ Senescence Green Flow Cytometry Assay Kit (Thermo Fisher, C10840) for SA-*β*-galactosidase activity. Cytokine concentrations in the matched supernatants were determined by ELISA for human IFN-*γ* (FANKEW, F0033-A) and human TNF-*α* (FANKEW, F0121-A) on an Epoch microplate reader (BioTek Instruments), with absolute concentrations extrapolated from the corresponding standard curves.

#### Elder validation cohort scRNA-seq

The out-of-distribution validation cohort was processed independently of the perturbation experiments above. Cryopreserved PBMC aliquots from elderly donors were rapidly thawed in a 37*^◦^*C water bath and transferred to 15-mL conical tubes containing 9 mL pre-warmed RPMI 1640 with 10% FBS. Suspensions were centrifuged at 300 *×* g for 5 min, washed twice in RPMI 1640 with 2% FBS, and counted by acridine orange / propidium iodide (AO/PI) staining. Because matched genotype variant-call-format (VCF) data were available, PBMCs from ten donors were pooled per single-cell capture reaction at equal viable-cell numbers, enabling genotype-based donor demultiplexing. To increase recovery under this multiplexed design, 80,000 viable cells were loaded per capture reaction, a pilot-optimised input for the SeekOne^r^ DD Single Cell 5’ Transcriptome V1.3 platform. Libraries were prepared per the manufacturer’s instructions and sequenced on the SURFSeq 5000 platform with a 29-bp Read 1 / 100-bp Read 2 strategy, targeting approximately 25,000 reads per cell.

### Benchmarking and statistical analysis

All statistical tests were two-sided unless stated. Multiple testing was corrected by Benjamini–Hochberg FDR. Donor was treated as a random effect in all mixed models. Hit-enrichment confidence intervals were computed by bootstrap (*n* = 10, 000). The primary claim of the RVQ–LLM model is improved phenotype-relevant prediction, not uniformly lower whole-transcriptome MSE everywhere. Data analysis used Scanpy v1.10.0, PyTorch v2.6.0, and custom Python v3.11.5 pipelines. Trained model weights, code, and agent logs are available as described in the Code availability statement.

## Supplementary information

The supplementary information accompanying this manuscript contains 16 supplementary figures paired with the main figures, plus supplementary tables.

## Acknowledgments

We thank Prof. Zeyu Chen for experimental advice.

## Author contributions

L.T., J.C. and X.L. conceived and supervised the study. Y.Y., J.K. and L.T. developed the RVQ tokenizer; G.L. built CellQ; Y.Y. and Y.F. constructed PACE and the agent. X.F., W.D., H.H., W.R. and S.L. performed the wet-lab perturbation screen, and X.F. performed the T-cell rejuvenation assay. G.H., J.S. and K.Z. acquired the Kunshan cohort. Y.Y. and L.T. wrote the manuscript with input from all authors.

## Competing interests

J.K. is a cofounder with Rectified Flow Capital Limited. Although not directly related to this paper, X.L. is a cofounder of iCamuno Biotherapeutics. The authors declare no other competing interests related to this work.

## Ethics approval and consent to participate

The KARE elderly validation cohort study was approved by the Independent Ethics Committee at the First People’s Hospital of Kunshan (protocol IEC-C-007-A07-V3.0). Cryopreserved PBMCs for perturbation experiments were obtained from STEMCELL Technologies, where all donor samples were collected under approved ethical protocols with documented informed consent. All experiments were performed in accordance with the Declaration of Helsinki.

## Consent for publication

Not applicable; no identifiable individual-level data are presented.

## Funding

This work was supported by the Major Project of Guangzhou National Laboratory (Grant Nos. GZNL2025A01006, GZNL2024A01015 and GZNL2024A03001).

## Data availability

The single-cell RNA-sequencing data for the 110-compound perturbation screen have been deposited in the Genome Sequence Archive (GSA) under accession code OMIX017117. The KARE elderly validation cohort data are available through the Kunshan Aging Research with E-health program under restricted access. Public datasets used for PACE construction are available at: Wang cohort (SYNAPSE, accession syn61609846, https://www.synapse.org), AIDA (CELLxGENE, https://cellxgene.cziscience.com/collections/ced320a1-29f3-47c1-a735-513c7084d508), and OneK1K (CELLxGENE, https://cellxgene.cziscience.com/collections/dde06e0f-ab3b-46be-96a2-a8082383c4a1). The Tahoe-100M perturbation atlas used for CellQ benchmarking is available at https://huggingface.co/datasets/tahoebio/Tahoe-100M.

## Code availability

All custom code for RVQ–LLM, CellQ, and PACE, the Analyzer–Planner–Auditor agent framework, and the trained model weights and agent logs are deposited on Zenodo (DOI). During peer review the record is set to restri tors and reviewers can access the files through the following share link: Upon acceptance the record will be made publicly accessible at the same DOI.

## Notes

### Competing Interest Statement

The authors have declared no competing interest.

